# Many but not all lineage-specific genes can be explained by homology detection failure

**DOI:** 10.1101/2020.02.27.968420

**Authors:** Caroline M. Weisman, Andrew W. Murray, Sean R. Eddy

## Abstract

Genes for which homologs can be detected only in a limited group of evolutionarily related species, called “lineage-specific genes,” are pervasive: essentially every lineage has them, and they often comprise a sizable fraction of the group’s total genes. Lineage-specific genes are often interpreted as “novel” genes, representing genetic novelty born anew within that lineage. Here, we develop a simple method to test an alternative null hypothesis: that lineage-specific genes do have homologs outside of the lineage that, even while evolving at a constant rate in a novelty-free manner, have merely become undetectable by search algorithms used to infer homology. We show that this null hypothesis is sufficient to explain the lack of detected homologs of a large number of lineage-specific genes in fungi and insects. However, we also find that a minority of lineage-specific genes in both clades are not well-explained by this novelty-free model. The method provides a simple way of identifying which lineage-specific genes call for special explanations beyond homology detection failure, highlighting them as interesting candidates for further study.

## Introduction

Homologs are genes that descend from a common evolutionary origin. “Lineage-specific genes” are defined operationally as genes that lack detectable homologs in all species outside of a monophyletic group [1]. Also referred to as “taxonomically-restricted genes” [2, 3], and as “orphan genes” when found only in a single species [4, 5], they are ubiquitous in the genomes of sequenced organisms. For example, by previous reports, 23% of *C. elegans* genes are specific to the *Caenorhabditis* genus [6]; 6% of honey bee genes are specific to insects [7]; 25% of ash tree genes are specific to the species [8]; and 1% of human genes are specific to primates [9].

Where do lineage-specific genes come from? A common interpretation is that they are “novel” genes. Various proposals for the molecular nature of this novelty have been advanced. For example, lineage-specific genes have been interpreted as “de novo genes” that have evolved from previously noncoding sequence [10, 11], and as duplicated genes that diverged radically in evolving a new function [12]. Though different in detail, these proposals share the key assumption that a lack of detectable homologs indicates some kind of biological novelty: lineage-specific genes either have no evolutionary homologs or no longer perform the same function as their homologs outside the lineage [13-16]. We refer to these interpretations collectively as the “novelty hypothesis” of lineage-specific genes. The novelty hypothesis has informed work on the evolution of new features at molecular, cellular, and organismal scales [16-20].

An alternative explanation for a lineage-specific gene is that nothing particularly special happened in the gene’s evolutionary history and homologs *do* exist outside the clade, but that computational similarity searches (e.g. BLAST) merely failed to find those homologs. We refer to such unsuccessful searches as homology detection failure. As homologs diverge in sequence from one another, the statistical significance of their similarity declines. Over evolutionary time, with a constant rate of sequence evolution, the degree of similarity may fall below the chosen significance threshold, resulting in a failure to detect the homolog. Some lineage-specific genes may just be those for which this happens to have occurred relatively quickly, even in the absence of any novelty-generating evolutionary mechanisms.

Here, we describe a method for evaluating whether this alternative hypothesis of homology detection failure is sufficient to account for a lineage-specific gene. We develop a mathematical model that estimates the probability that a homolog would be detected at a specified evolutionary distance if it was evolving at a constant rate under standard, novelty-free evolutionary processes. We apply the method to lineage-specific genes in insects and yeasts, and find that many, but not all, lineage-specific genes in these taxa can be explained by homology detection failure.

## Results

### A null model of homolog detectability decline as a function of evolutionary distance

We developed a formal test of the null hypothesis that homology detection failure is sufficient to explain the lineage-specificity of a gene. Specifically, we model the scenario in which the gene actually existed in a deeper common ancestor, evolved at a constant rate, and has homologs outside the clade in which its homologs are detected that appear to be absent solely due to homology detection failure. This is an evolutionary null model: it invokes no processes beyond the simple scenario of orthologs diverging from a common ancestor and evolving at a constant rate.

Because of its use in previous work on lineage-specific genes and in sequence analysis more broadly, we use BLASTP as the search program used to detect homologs here. In search programs like BLAST, sequence similarity is used to infer homology between two genes. Such programs report a similarity score (referred to as “bitscore” by BLAST) between a pair of sequences, as well as the number of sequences that would be expected to achieve that similarity score by chance (an E value). When this number falls below a significance threshold (e.g. E < 0.001), statistically significant similarity is interpreted as evidence that the two genes are homologous. The similarity score therefore directly determines whether a homolog is successfully detected in a search.

The key idea in our method is to predict how the similarity score between two homologs evolving according to our null model is expected to decline as a function of the evolutionary distance between them. We can then ask whether a given gene’s lack of detectable homologs outside of the lineage is expected under this null evolutionary model. Previous work on this problem has simulated the evolution of each gene [21-24]. In preliminary work, we explored similar ideas but found results from detailed simulations to be fragile. A simulation approach requires selection of many evolutionary parameters, which has led to questions about the sensitivity of results to these details [23-27].

We chose instead to analytically model how the similarity score between two homologs decays with the evolutionary distance between them. Briefly, our model assumes that the similarity score between two homologs is proportional to the percent identity between them, and that every position in the protein mutates at the same protein-specific rate, which is constant over evolutionary time. With these assumptions, the expected similarity score *S* between two homologs separated by an evolutionary divergence time *t* is given by *S(t)* = *a*e^-*bt*^, where the protein-specific parameters *a* and *b* are related to the protein’s length and the protein’s evolutionary rate, respectively. The variance of this similarity score is given by σ^2^ = *a*(1-e^-*bt*^)(e^-*bt*^). A fuller explanation of these formulas can be found in the methods.

We can predict similarity scores for a given gene if we have three inputs: a gene from a chosen focal species, the similarity scores of successfully identified homologs of the gene in a few other species *(S)*, and the evolutionary distances between the focal species and these other species *(t).* (As described in the following section and methods, we precalculate these evolutionary distances *t* from an aggregate of many genes from the set of species under consideration, and therefore they do not depend on the particular gene under consideration.) We use these inputs to find the gene-specific values of the parameters *a* and *b* that produce the best fit to our equation describing how similarity scores decline within the species where homologs were detected. We then use these parameters to extrapolate and predict the expected similarity score of hypothetical homologs of the gene at evolutionary distances beyond those of the species whose homologs were used in the parameter fitting. Given an E-value threshold, this predicted similarity score, and the expected variance of the similarity score, we can estimate the probability that a homolog will be undetected at these longer evolutionary distances. In the analyses that follow, we use a relatively permissive E-value threshold of 0.001.

This key idea is illustrated in Figure 1, which shows examples of fitting similarity scores versus evolutionary distance for several different yeast and insect genes.

**Figure 1:**
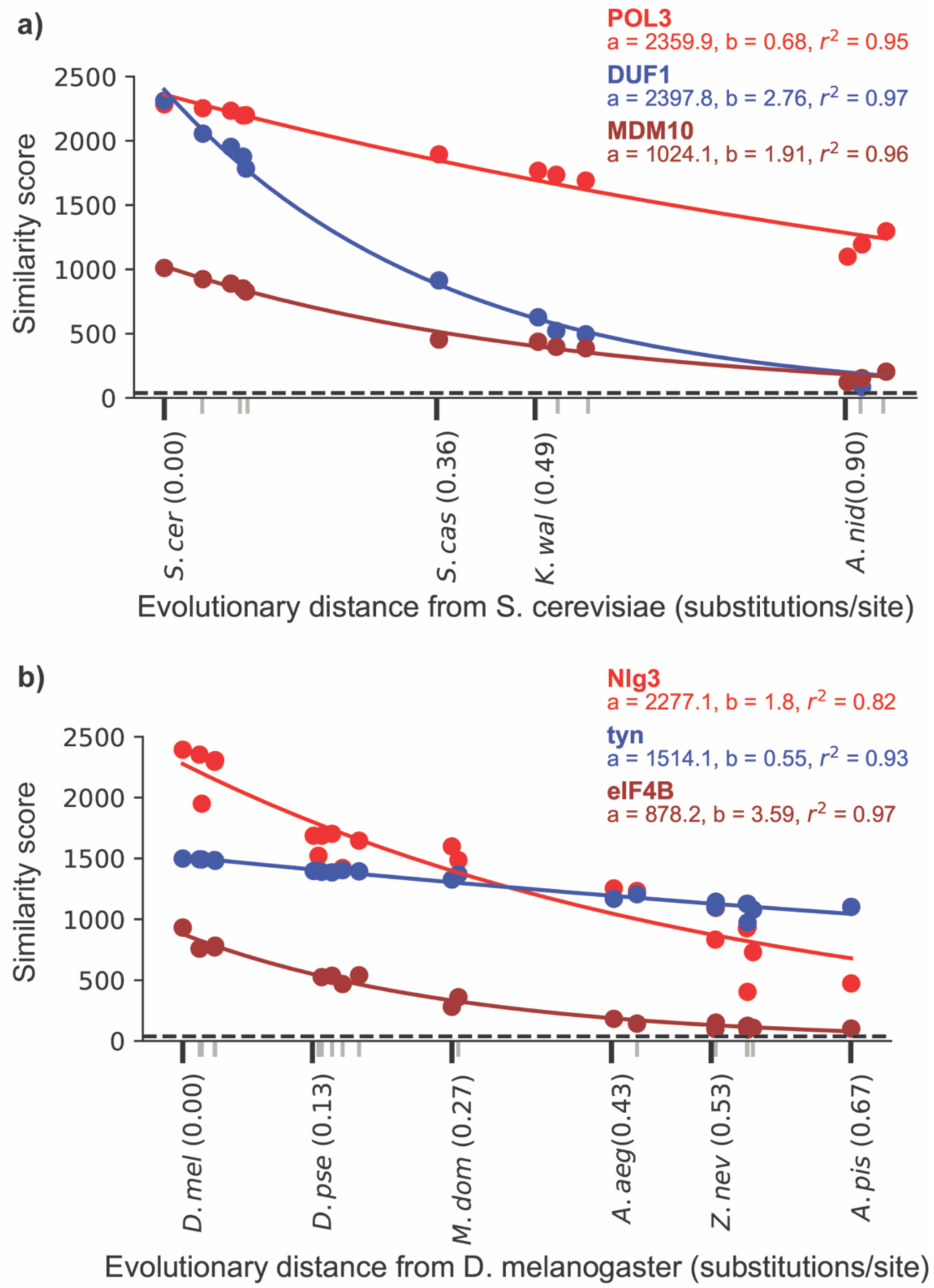
Depictions of the fit of the null model of similarity score decline with evolutionary distance for three representative proteins from *S. cerevisiae* (a) and *D. melanogaster* (b). Colored points represent the BLASTP score between the protein and its ortholog in the species that is at the evolutionary distance indicated on the x axis. Tick marks on the x axis represent each of the species used here. For visual clarity, only some species names and evolutionary distances are included, indicated with black tick marks; gray tick marks represent the other unlabeled species. The dashed line represents the detectability threshold, the score below which an ortholog would be undetected at our chosen E-value of 0.001. The best fit values of *a* and *b* are shown for each protein. The r^2^ value is also shown and was calculated from a linear regression of the log of the similarity score versus evolutionary distance.

### The null model adequately describes the decay of ortholog detectability with evolutionary distance

We applied our model to genes of the yeast *S. cerevisiae* and the fly *D. melanogaster* and their orthologs in several fungal and insect outgroups respectively. We focus on the fungi and insects because their genomes are well-annotated, they have closely related and well-annotated sister species, and they have been the focus of previous work on lineage-specific genes [5, 11, 28-31]. For *S. cerevisiae*, we included 11 fungal species spanning a divergence time of ∼600 million years [32]; for *D. melanogaster*, we included 21 insect species spanning a divergence time of ∼400 million years [33]. These species are listed in Figure 2.

**Figure 2:**
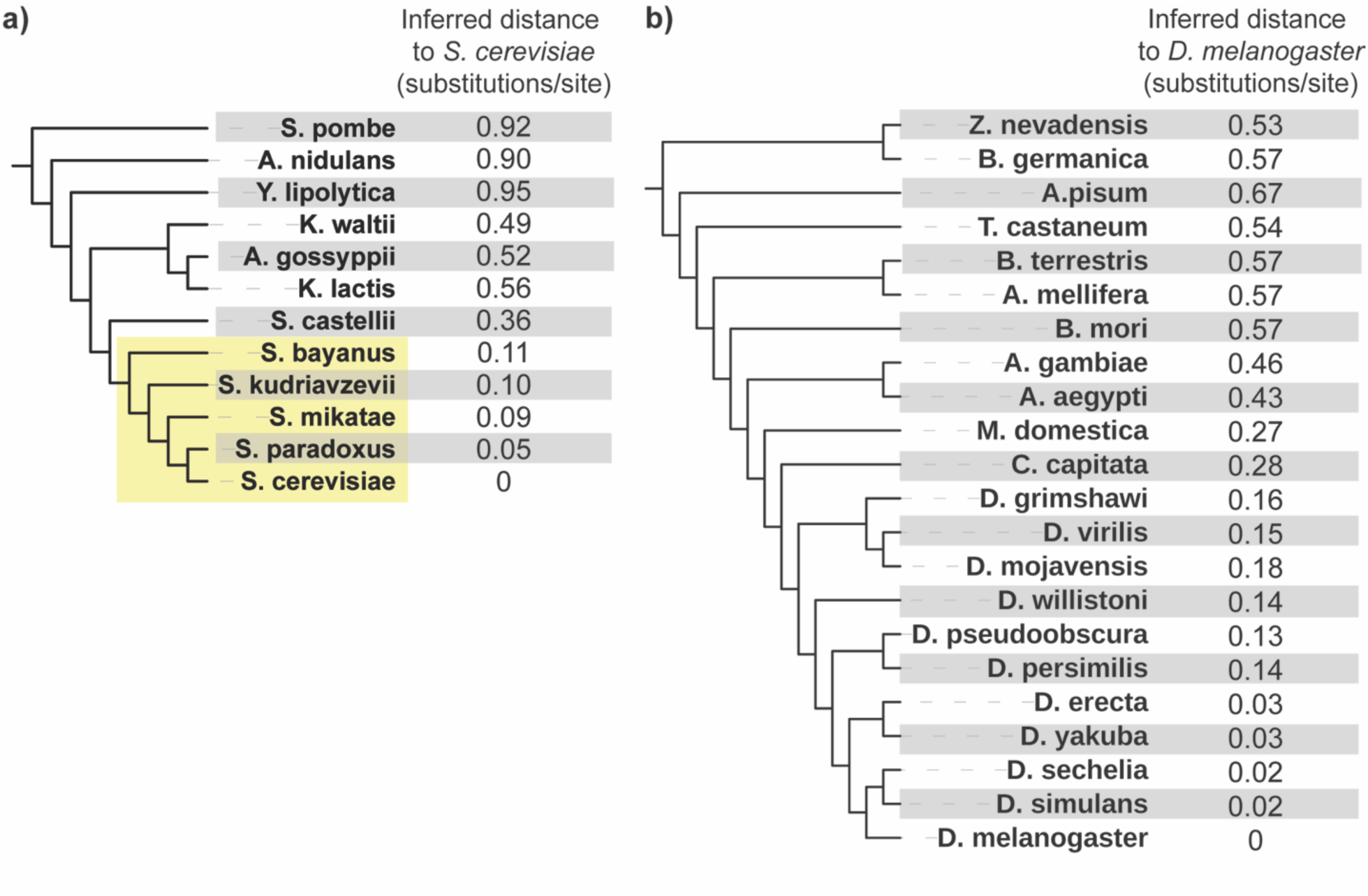
Inferred evolutionary distances between each fungal species and *S. cerevisiae* (a) and each insect species and *D. melanogaster* (b). The tree topologies for these taxa are based on previously published studies [33, 35] and were not calculated here; branch lengths are not to scale. The fungal *sensu stricto* lineage, referenced frequently in the text, is shaded in yellow.

Before using our null model to ask whether it explains the lack of detected homologs of lineage-specific genes, we confirmed that it is a good approximation of how similarity scores decay with evolutionary distance. To do this, we tested how well the model represents the decay of similarity scores of general *S. cerevisiae* and *D. melanogaster* genes in increasingly distant species. If the model fits this decay well for most genes, it is likely a good representation of the minimal evolutionary process in the null hypothesis, and can therefore detect deviations from that process.

To obtain evolutionary distances from the focal species (*t* values), represented by the x axis in Figure 1, we used 102 genes from the Benchmarking Universal Single Copy Ortholog (“BUSCO”) [34] database to calculate evolutionary distances in substitutions/site between *S. cerevisiae* and each of the 11 other fungi, and 125 BUSCO genes to calculate evolutionary distances between *D.* melanogaster and each of the 21 other insects (Methods). In both taxa, to show that distances can be reliably computed using a small number of genes, we also re-calculated these distances using two random subsets of 15 BUSCO genes. Distances computed from these different gene sets were similar (Supplemental Table 1). Figure 2 shows evolutionary distances inferred from one of the 15 gene sets between the focal organism *S. cerevisiae* and the 11 other fungi, and between the focal organism *D. melanogaster* and the 21 other insects. For reference, Figure 2 depicts these distances along with a topology taken from previous phylogenetic studies of these taxa [33, 35]; branch lengths are not to scale. We use these distances, computed from one of the 15 gene subsets, in all results presented in the main text below.

We next took all annotated *S. cerevisiae* and *D. melanogaster* proteins (Supplemental Table 2) and identified the similarity scores of their detectable orthologs in each of the 11 other fungal and 21 other insect outgroup species respectively. (For *S. cerevisiae* and *D. melanogaster*, the score is the comparison of the protein with itself.) We identified orthologs using reciprocal best BLASTP search with a threshold of E < 0.001 (Methods). Reciprocal best BLASTP is not a perfect means of distinguishing orthologs from paralogs, and results in some genes failing to be assigned to orthologs in some species, but suffices for the purpose and is easy to do at scale.

With these similarity scores (*S*) and evolutionary distances (*t*) in hand, we tested how well our model explains the observed decline in similarity scores with increasing evolutionary distance in fungal and insect genes. Our model predicts a linear relationship between the log of ortholog similarity scores and evolutionary distance. We therefore assessed the fit of the model by performing a linear regression of the log of each protein’s similarity score, ln *S(t*), against the inferred evolutionary distance to the focal species, *t*, and computing the square of the Pearson correlation coefficient (r^2^), which measures how much of the variance in ln *S(t*) is explained by *t*.

The model predicts similarity scores reasonably well. The mean and median r^2^ were 0.92 and 0.95 for similarity scores of *S. cerevisiae* genes (Supplemental Figure 1b). We repeated this with *D. melanogaster* proteins and their orthologs in the other insects, where the mean and median r^2^ were 0.84 and 0.91 for similarity scores of *D. melanogaster* genes (Supplemental Figure 1e). Results were similar using the two other sets of estimated distances (Supplemental Figure 1).

As well as considering the fit of each gene to the expected value of the model, we tested how well our estimate for the variance of the similarity score captured the observed scatter around this expected value. To do this, for the ortholog of each *S. cerevisiae* gene in each species, we calculated the difference between the actual and expected similarity score and expressed it as a multiple of the predicted standard deviation σ = √*a*(1-e^-*bt*^)(e^-*bt*^) of the similarity score (a Z score). We expect these Z-scores to follow a normal distribution if our model’s estimated variance is correct, which is roughly what we observe (Supplemental Figure 2a-c). Approximately 92% of *S. cerevisiae* orthologs have observed scores within 3 s.d. of the prediction; for a standard normal distribution, 99% are expected. 7% of scores are below three s.d., and 1% are above three s.d. Results in *D. melanogaster* are similar: 88% have observed scores within 3 s.d., 8% below, and 4% above. We attribute the skew toward predicted scores that are higher than observed scores to the fact that our model neglects how insertions and deletions may disrupt the length of a local alignment. Results were similar when using the two other sets of estimated distances (Supplemental Figure 2d-f).

We asked whether the best-fit values of the parameters *a* and *b* found for the fungal proteins are correlated with the interpretation of these parameters in our model. We expect values of *a* to be related to gene length, and values of *b* to be related to evolutionary rate. Using comparisons to *S. cerevisiae* genes, we plotted *a* versus gene length and *b* vs. maximum likelihood estimates of evolutionary distance in substitutions/site in multiple alignments of proteins from *S. cerevisiae* and the four most closely related species. The *a* parameter is indeed highly correlated with gene length (r^2^ = 0.99), and *b* is more weakly correlated with gene-specific evolutionary rate (r^2^ = 0.47) (Supplemental Figure 3). The distributions of the estimated *a* and *b* parameters across all genes are long-tailed and approximately log-normal (Supplemental Figure 4), consistent with other analyses of distributions of gene length [36] and evolutionary rate [37].

### Many lineage-specific genes can be explained by homology detection failure

Having validated our null model for similarity score decline, we then focused on lineage-specific genes and used the model to ask our central question: how often is homology detection failure alone enough to explain a lineage-specific gene?

We first considered annotated *S. cerevisiae* proteins that are lineage-specific to the *sensu stricto* yeasts, a young lineage sharing a common ancestor ∼20 Mya containing the five species *S. cerevisiae, S. paradoxus, S. mikatae, S. bayanus, and S. uvarum* (Figure 2a), which has been the focus of previous work on lineage-specific genes [11, 30]. We identified 375 such *sensu stricto-*specific genes, defined as having homologs detectable by BLASTP in at least one of these species but lacking detectable homologs in the nearest outgroup *S. castellii* or in any other outgroups according to a permissive E-value threshold of 0.001 (Methods). Between 40 and 70% of *sensu stricto* specific genes identified in two previous studies are included in this set [11, 30]. The remainder are either ORFs not used in our initial search because they are marked as dubious in both the Saccharomyces Genome Database and Refseq and so have been removed from the *S. cerevisiae* Refseq annotation, or because we detected homologs outside of the *sensu strictos*, likely due to our permissive E-value threshold. Since our detectability model is regression-based, we only use it on genes with a minimum of 3 observed homologs (including the gene in the focal species); for example, we could not perform this computation on the *S. cerevisiae* gene *BSC4* [38], proposed to have a very recent de novo origin and thus only found in *S. cerevisiae*. We applied our model to 155 *sensu stricto*-specific proteins.

For each of these 155 lineage-specific genes, we used the best-fit values of the *a* and *b* parameters found above to extrapolate and predict the score of an ortholog at the evolutionary distance of *S. castellii* under the null model. Using parameters from the *sensu stricto* lineage to extrapolate to more distant species corresponds to assuming that these two groups of orthologs have evolved in the same manner since their divergence from their common ancestor. Finally, we calculated the probability that a homolog at the evolutionary distance of *S. castellii* would be detected, P(detected | null model, t_*castellii*_), by using our model for similarity score variance to generate a probability distribution for the score and computing the percentage of the probability mass in this distribution below our chosen detectability threshold (corresponding to an E-value of 0.001).

This analysis is illustrated for one example of a *sensu stricto*-restricted *S. cerevisiae* protein, Uli1, in Figure 3. Uli1 has been implicated in the unfolded protein response [39], making it one of only a few *sensu stricto* specific genes with experimental evidence of function, and its lineage-specificity has prompted previous studies to propose that it originated de novo [11, 30]. However, we find that the probability that an ortholog of this gene would be detectable in *S. castellii*, P(detected | null model, t_*castellii*_), is approximately 0, indicating that a null evolutionary model is sufficient to explain the lineage specificity of this short and rapidly-evolving gene.

**Figure 3:**
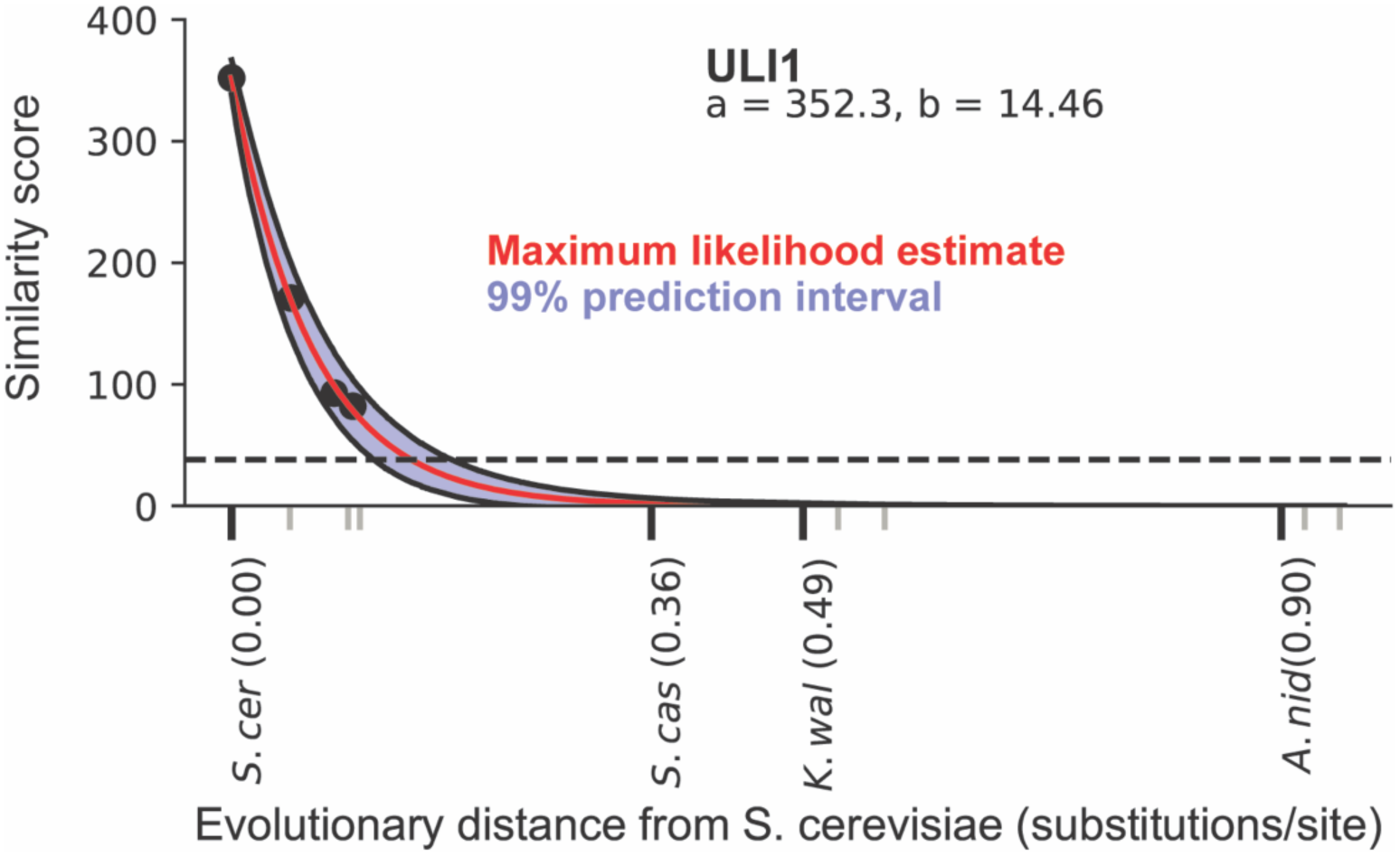
Illustration of the prediction of detectability decline for the *S. cerevisiae* protein Uli1, displayed as in Figure 1. At the evolutionary distance of the nearest outgroup *S. castellii*, the entire prediction interval lies below the detectability threshold, indicating a ∼0% probability that an ortholog would be detected under the null model even if an *S. castellii* ortholog were present.

The result of performing this test on all of these 155 *sensu stricto*-specific genes is shown in Figure 4a, which depicts the distribution of probabilities of detecting a homolog in the outgroup *S. castellii* given the null model and the evolutionary distance between *S. cerevisiae* and *S. castelli*, P(detected | null model, t_*castellii*_). Many genes have a very high probability of being undetected, and a majority are more likely to be undetected than detected: 55% have P(detected | null model, t_*castellii*_) below 0.05, and 73% percent below 0.5. This implies that homology detection failure is sufficient to explain a large number, potentially a majority, of these lineage-specific genes. Homologs of these genes only being detected in *sensu stricto* species does not require invoking evolutionary novelty.

**Figure 4:**
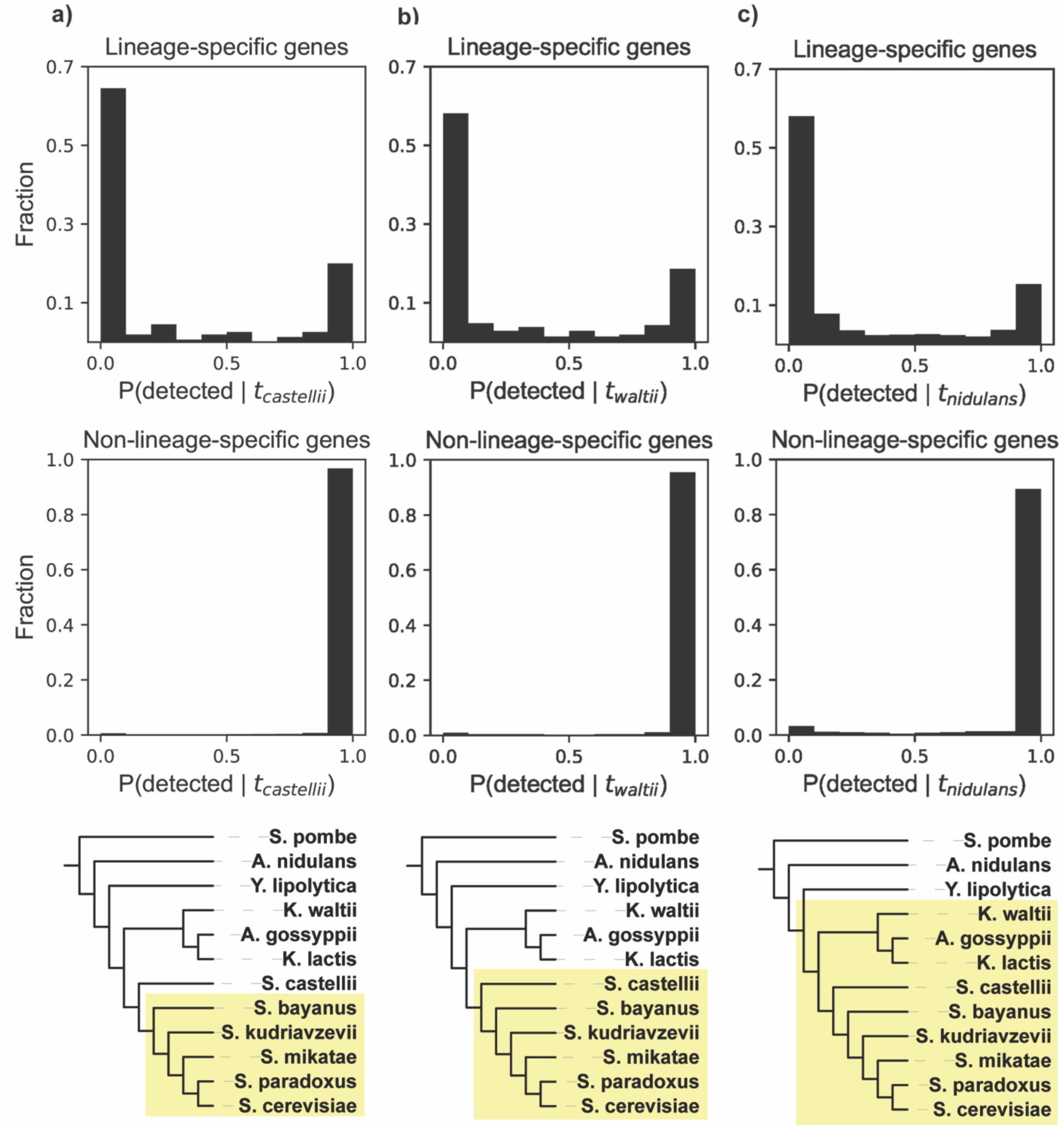
Distributions of detectability prediction results for three yeast lineages (a, b, c). Top: results for all lineage specific genes. Middle: results for all non-lineage specific genes. Bottom: lineage tested as shown by yellow shading. In c), note that *Y. lipolytica* is the topological outgroup to the shaded lineage, but is not the closest species by evolutionary distance (branch lengths are not to scale).

We repeated this procedure for *D. melanogaster* genes restricted to the *Drosophila* genus. This young lineage shared a common ancestor ∼70 Mya, with the housefly *M. domestica* as the nearest outgroup in our analyses (Figure 5a). We identified 1611 *Drosophila-*restricted genes (Methods), of which 1273 had enough identified orthologs in the *Drosophila* lineage to perform our analysis. Again, many *Drosophila*-restricted genes are very likely to be undetected: 46% percent have values of P(detected | null model, t_domestica_) below 0.05, and 76% percent are below 0.5 (Figure 5a). Homology detection failure is therefore also sufficient to explain many lineage-specific genes in this group.

**Figure 5:**
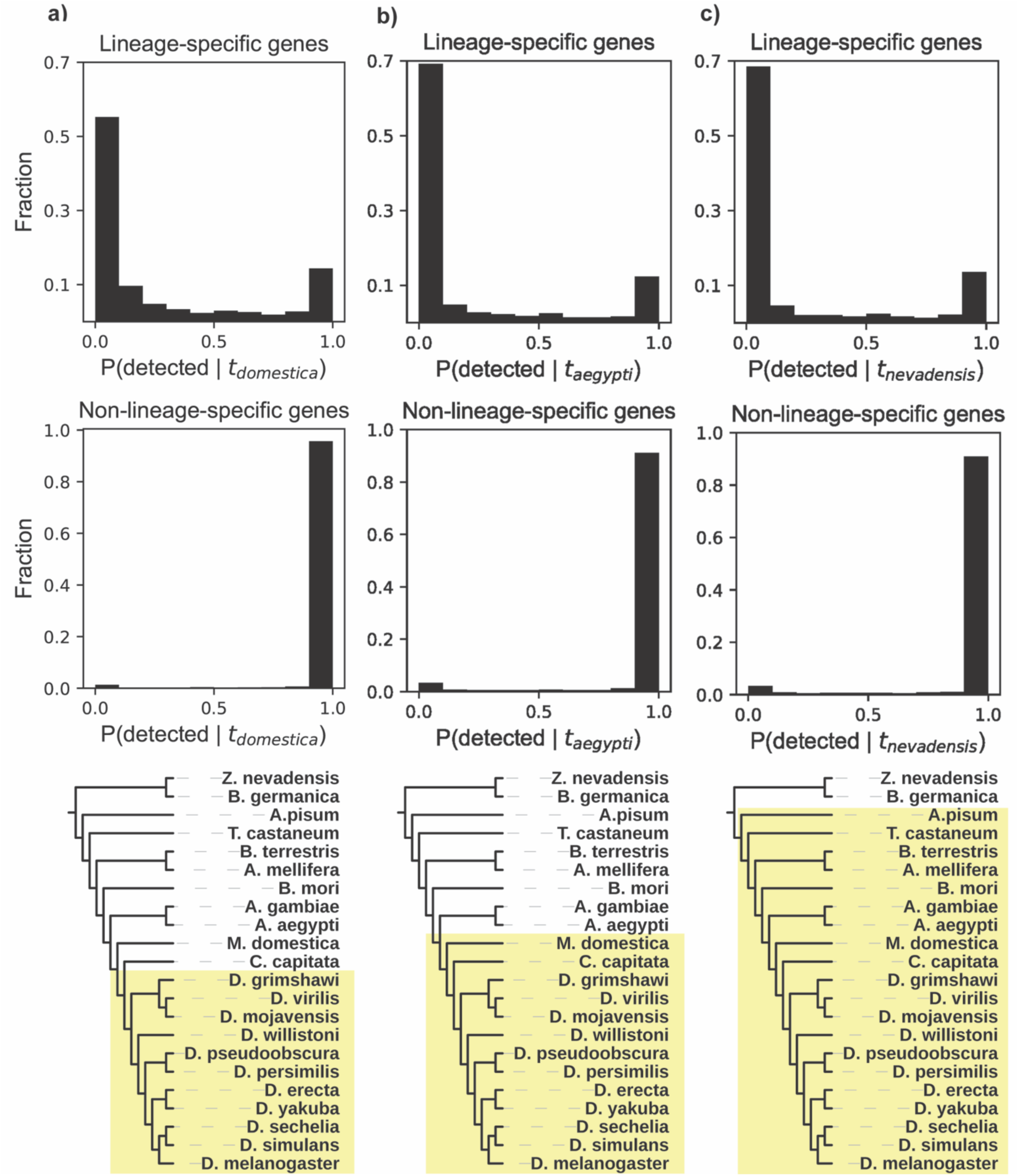
Distributions of detectability prediction results for three insect lineages (a, b, c). Top: results for all lineage specific genes. Middle: results for all non-lineage specific genes. Bottom: lineage tested as shown by the yellow shading. In a), note that *C. capitata* is the topological outgroup to the shaded lineage, but is not the closest species by evolutionary distance (branch lengths are not to scale).

As both the *sensu stricto* yeasts and the drosophilid flies are relatively young lineages, we asked whether these results generalize to older lineages. In fungi, we tested two additional lineages with approximate divergence times of ∼70 Mya (Figure 4b) and ∼250 Mya [32] (Figure 4c). In insects, we also tested two additional lineages, with approximate divergence times of ∼150 Mya (Figure 5b) and ∼350 Mya [33] (Figure 5c). We calculated P(detected | null model, t_outgroup_) for each of these four additional lineages. Results in all of these lineages are very similar to those in the younger lineages tested above: we predict that a large number of lineage-specific genes have very low probabilities of being detected, with a majority more likely to be undetected than detected (Figures 4b,c, 5b, c). Homology detection failure is thus sufficient to explain a large number of lineage-specific genes in these older lineages as well.

As a control, we asked our model to predict the probability of detecting homologs of genes that are *not* lineage-specific, meaning that these genes have homologs that are detected both inside and outside of the lineage. We repeated the same procedure on all non-lineage-specific genes in the six lineages tested above. As we did for the lineage-restricted genes, we used only similarity scores from orthologs within the given lineage to calculate the probability of detecting homologs in the nearest outgroup to the lineage, P(detected | null model, t_outgroup_). If our model operates correctly, it should predict high values of P(detected | null model, t_outgroup_) for these genes, since their homologs are in fact detected. In accordance with this expectation, our model predicts that the vast majority (>97% in all lineages) of these genes have a very high probability of being detected, P(detected) > 0.95 (Figures 4, 5). This analysis, like earlier analyses, was robust to the use of different sets of genes for calculating evolutionary distances (Supplemental Table 3).

### More sensitive homology searches detect beyond-lineage homologs for many lineage-specific genes well-explained by homology detection failure

If a gene being lineage-specific is due to the failure of BLASTP to detect homologs that are in fact present, we would expect that a more sensitive search will sometimes succeed in finding homologs where BLASTP did not. We asked whether this was the case for genes whose lineage-specificity was consistent with the hypothesis of detection failure: can we use a more sensitive method to find previously undetected homologs for these genes? We refer to such homologs, detected using a different method in species outside of the originally defined lineage, as “beyond-lineage homologs.”

We used *sensu stricto* yeast-specific genes as a case study to ask this question. These yeasts and several of their nearest outgroups have a high degree of conservation of chromosomal gene order (synteny), presenting the opportunity for a more sensitive search. A standard similarity search tests all proteins in a large database of sequences, such as a complete proteome. The resulting multiple testing burden requires a higher score to achieve statistical significance than would be required for a search over a smaller number of sequences. In these yeasts, synteny allows us to restrict a similarity search to one candidate gene at the orthologous chromosomal locus, reducing the multiple testing burden and enabling ortholog identification with a lower score. For the fungal species used here, a proteome-wide search would need a BLASTP score of ∼37 to achieve an E-value of 0.001, but a single-protein search would only require a score of ∼24. Orthologs with scores between these two values would be missed in our initial search but successfully detected with synteny-guided similarity searches.

We used this strategy to search for beyond-lineage orthologs for all *sensu stricto-*specific genes for which the null model of detection failure is a reasonable explanation. We use a threshold of P(detected | null model) < 0.95 to define these genes. This choice is a conservative threshold that corresponds to genes that are insignificant according to a traditional significance test threshold of P(undetected | null model) = 1 – P(detected | null model) > 0.05. There are 126 *sensu stricto*-specific genes that pass this threshold.

To identify the orthologous locus in outgroup yeasts for these 126 *S. cerevisiae* genes, we used the Yeast Gene Order Browser (YGOB), an online resource that curates the chromosomal orthology relationships between species including the *sensu stricto* yeasts, *S. castellii, K. waltii, A. gossypii*, and *K. lactis* [40]. 24 of these 126 *sensu stricto*-specific genes have an orthologous locus in at least one of these outgroup yeasts listed in YGOB. For all of these genes, the upper bound of the 99% prediction interval for the similiarity score predicted by our model is above the detectability threshold of 24 bits, indicating that they are potentially detectable by this analysis. Of these 24 genes, 11 had an annotated gene at the orthologous locus in at least one outgroup species with significant (E<0.001) similarity to the *S. cerevisiae* gene. In all but 2 of these cases, the similarity score fell within our prediction interval (in those 2 cases, the similarity score was slightly higher than predicted). These 11 genes and their proposed orthologs are listed in Supplemental Table 4.

In total, we found beyond-lineage homologs for 46% of genes for which we were able to perform a synteny analysis. We note that this is a conservative estimate. We only considered ORFs that are already annotated in outgroup species, although unannotated orthologs may be present. Additionally, the lower bound of the 99% similarity score prediction interval for all remaining 54% of these genes is lower than the threshold required for detection via synteny, so that all have some probability of orthologs still being missed in this analysis.

### Some lineage-specific genes are poorly explained by homology detection failure

In all lineages studied here, there are also lineage-specific genes that are poorly explained by the null hypothesis: their similarity score declines too slowly to make homology detection failure alone a good explanation for their lineage-specificity. These are the genes with high values of P(detected | null model). In all six lineages we studied, 10-20% of lineage-specific genes have detection probabilities of 0.95 or greater (Figures 4,5).

This result is illustrated by one such *sensu stricto-*specific protein, Spo13, in Figure 6. Spo13 has been proposed as a candidate de novo gene [30] by virtue of its lineage-specificity, and this analysis highlights it as a particularly promising novel gene candidate amongst the large number of other lineage-specific genes in the *sensu stricto* lineage.

**Figure 6:**
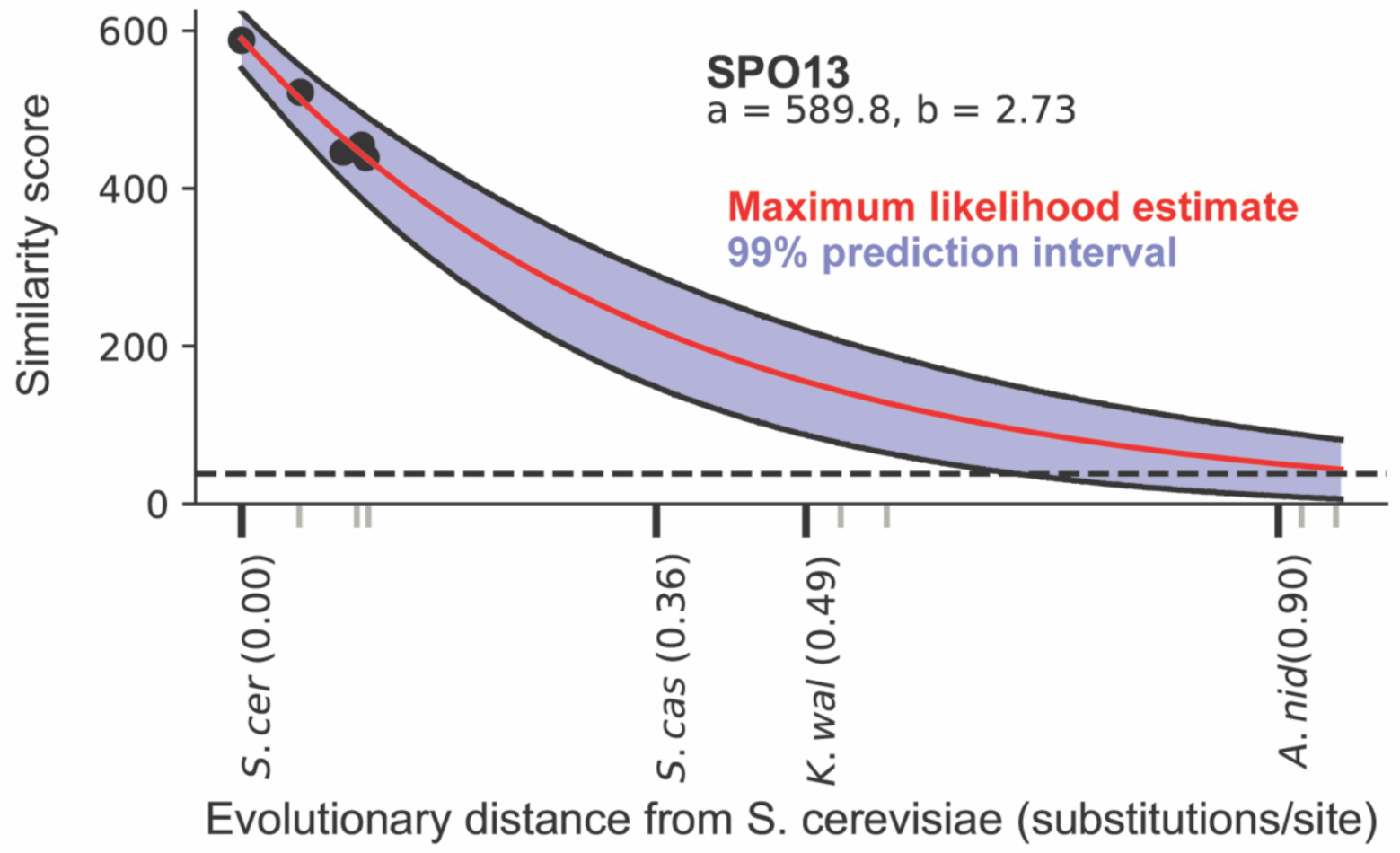
Detectability prediction results for the *S. cerevisiae* protein Spo13, displayed as described in Figure 3. At the evolutionary distance of the nearest outgroup *S. castellii*, the entire prediction interval lies well above the detectability threshold, indicating a ∼100% probability that an ortholog should be detected in this species under the null model.

The existence of lineage-specific genes like Spo13, which our null model predicts should have detectable homologs outside of the lineage, indicates that evolutionary mechanisms beyond those included in the null model may be operating. Among such mechanisms are those postulated by the novelty hypothesis, like de novo origination and duplication-induced neofunctionalization. However, other known mechanisms could also explain such genes. These include processes that cause the gene tree to deviate from the species tree, like horizontal gene transfer and any mechanisms that change the evolutionary rate of a protein on a restricted part of the tree.

### Characterization of yeast lineage-specific genes that are poorly explained by homology detection failure

We next aimed to characterize genes whose lineage-specificity is poorly explained by homology detection failure. We again used *sensu stricto*-specific genes as a case study, allowing for synteny analysis and the biological insight provided by many genes in *S. cerevisiae* being comparatively well-studied. We selected the subset of *sensu stricto*-specific genes, including Spo13, whose lineage-specificity is poorly explained by homology detection failure, i.e. for which P(detected | null model) > 0.95. These are genes on the other side of the threshold applied above: the null hypothesis strongly predicts that homologs should be detected, making their lineage-specificity incompatible with the null hypothesis. There are 25 *sensu stricto*-specific genes that satisfy this threshold. While a thorough study of these genes is beyond our scope, we report a few initial observations.

“De novo origination,” the process of a new gene emerging from previously non-coding sequence, is a commonly proposed origin of lineage-specific genes [12]. We asked how many of these 25 lineage-specific genes could plausibly be such de novo genes. By definition, genes that have emerged de novo in the *sensu stricto* lineage should have no out-of-lineage homologs, and so the more sensitive synteny-based homology search strategy used above should fail to find such homologs. We performed a synteny-based search for out-of-lineage homologs for these 25 genes in the same way as above. For 20 of these 25 genes, an orthologous locus is listed in YGOB. Of these 20, 12 have annotated genes with significant similarity (E<0.001) at the orthologous locus in at least one outgroup species. Thus, 12 of 25 genes, or just under half, of genes that are not well-explained by homology detection failure did not originate de novo in the *sensu stricto* lineage. This is a conservative estimate of the total number of genes that have out-of-lineage homologs, since, as described above, even this synteny-based homology search has finite sensitivity. Spo13, the gene shown in Figure 5, is one example of these lineage-specific genes that nonetheless are not de novo originated: it has out-of-lineage orthologs identifiable by synteny in *S. castellii, K. waltii, K. lactis*, and *A. gossypii*.

Genes that acquire a new function following duplication and divergence (“neofunctionalization”) are another proposed source of lineage-specific genes [12]. We therefore asked how many of our *sensu stricto*-specific genes have a paralog, consistent with the hypothesis that they emerged through duplication and divergence. Based on BLASTP searches within the *S. cerevisiae* genome, we find that 4 of the 25 lineage-specific genes have annotated paralogs specific to some subset of the *sensu stricto* yeasts, which therefore likely emerged after their divergence from *S. castellii*. We also find using YGOB that another 4 of these 25 genes have annotated paralogs resulting from the yeast whole genome duplication, which occurred before the divergence of *S. castellii* from the *sensu stricto* yeasts. In total, 8/25, or fewer than one-third, of these genes show evidence of having been the result of duplication events. However, we note that this estimate for the number of genes with paralogs is again conservative due to the finite sensitivity of the homology searches.

Finally, we performed a gene ontology enrichment test (Methods) to determine if certain biological processes were statistically overrepresented among these 25 genes. We find significant enrichment of genes involved in several GO categories relating to spore formation and meiosis, including “ascospore-type prospore membrane assembly” (p = 7*10^−5^; 3 observed vs 0.7 expected) and “meiotic cell cycle process” (p = 5*10^−5^; 7 observed vs 1 expected). Spo13, involved in meiotic cell cycle regulation through its roles in maintaining sister chromatid cohesion during meiosis I and promoting kinetochore attachment [41], is one such example. A table of these 25 genes and the features discussed above can be found in Supplemental Table 5.

## Discussion

The widespread interpretation of lineage-specific genes as evolutionarily novel assumes that absence of evidence for detectable homologs in outgroups is evidence that homologs are absent. The model we have presented here allows us to formally test the alternative, null hypothesis: homologs do exist outside the specified lineage, but they have diverged, at a constant novelty-free evolutionary rate, beyond the ability of a similarity search program to detect them. We find that this hypothesis is sufficient to explain a large number of lineage-specific genes in two taxa where lineage-specific genes have been interpreted as exhibiting some kind of evolutionary novelty.

Our results caution against automatically assuming that lineage-specific genes are novel. We cannot exclude the possibility that some genes explainable by homology detection failure are nonetheless novel: failing to reject a null hypothesis is not the same as accepting it. However, we argue that in the absence of additional evidence, the more conservative hypothesis of detection failure should be preferred to the more exotic one of evolutionary novelty. Our case study in the *sensu stricto* yeasts finds that more sensitive synteny-based homology searches successfully find previously undetected homologs for many lineage-specific genes, supporting this preference.

Although we find that many lineage-specific genes can be adequately explained by homology detection failure, we also find a minority of lineage-specific genes in fungi and insects that cannot. This leaves open the possibility that these genes are biologically novel. However, the reason that these genes reject our null model is not addressed by our present work. Our initial analyses do show that many of these genes are neither de novo genes nor have detectable paralogs, suggesting that processes other than the commonly proposed hypotheses of de novo origination and duplication-divergence may be at play. There are many possible processes that could cause genes to deviate from our null model, but one speculative example lies in the observed enrichment in yeast of genes involved in meiotic processes, exemplified by Spo13. This strikes us as suggestive of meiotic drive phenomena, which have been observed in yeast [42] and have been shown to cause rapid protein divergence [43], producing clade-specific rate accelerations leading to lineage-specific genes. More detailed characterization of these genes is required to understand if and in what way they are evolutionarily novel.

Another recent paper used a different approach to estimate the fraction of lineage-specific genes that are attributable to homology detection failure [44]. Vakirlis et al. used a small set of “microsyntenic blocks” to count how often a gene is not recognizably similar to its presumptive homolog in the syntenic position in a comparative genome. Assuming this sample approximates the frequency at which homologous genes diverge beyond recognition, they extrapolate this fraction to all genes. They conclude that 20-45% of lineage-specific genes in yeast, fly, and human phylogenies are attributable to homology detection failure. We consider this result to be in broad agreement with ours, although our results suggest somewhat larger estimates, and future effort will be needed to reconcile the two results to say whether homology detection failure accounts for a majority or a minority of lineage-specific genes.

There is increasing consensus that homology detection failure is frequent. It should be taken into account in studies that aim to use lineage-specific genes to identify candidates for genetic novelty. Our approach allows us to determine whether a *particular* lineage-specific gene is attributable to homology detection failure, and our approach is generalizable to a wide range of taxa, beyond the well-studied clades where synteny analysis can be used. We expect it to be useful in the wide variety of studies that aim to identify “new” genes that may underlie the evolution of morphological, behavioral, and other novel traits [7, 45-49]. An implementation of our method is freely available as source code at github.com/caraweisman/abSENSE, and as a web server at eddylab.org/abSENSE.

## Methods

### Model of similarity score as a function of evolutionary distance

Our model assumes that a similarity score *S* between two homologs is proportional to the number of sites that are identical in these two homologs. After the homologs diverge from their common ancestor, we assume that they undergo a substitution-only mutation process, in which each site in the proteins mutates into a non-identical site at the same protein-specific rate *b* per unit of evolutionary time. Neglecting the possibility of reversion, the probability that a site will *not* undergo a mutation within a time *t* following homolog divergence is *e*^*-bt*^, as given by the Poisson distribution. Given a constant number *N*_*0*_ of total sites in the protein, the number of sites that remain identical at that time *t* is binomially distributed with mean *N*_*0*_*e*^*-bt*^. If each identical site contributes the same amount *c* to the total similarity score, then the mean similarity score at time t is *S(t)* = c*N*_*0*_*e*^*-bt*^. The variance of the similarity score at time t is σ^2^ = c*N*_*0*_(1-e^-*bt*^)(e^-*bt*^), from the variance of a binomial distribution. In the text, we refer to c*N*_*0*_ as *a*, as these two parameters only appear as a product.

The simplifying approximations in this model abstract away the detailed effects of the substitution score matrix, insertion and deletion scores, and local versus global sequence alignment. Nonetheless, they appear to suffice to approximate *S(t)* in the regime of observable (i.e. statistically significant) local alignment scores *S(t)* >> 0, although we somewhat underestimate the empirically observed variance, and have some skew toward overestimating *S(t)* (Supplemental Figure 2).

### Identification of *S. cerevisiae* and *D. melanogaster* orthologs

We downloaded previously annotated proteomes of *S. cerevisiae, D. melanogaster*, and the other yeast and insect species indicated in Table 1 from several sources, largely Refseq and GenBank. Accession IDs for Refseq and GenBank proteomes and download links for those from other sources are listed in Supplemental Table 2. We performed a BLASTP (version 2.8.0) search [50] with an E-value threshold of 0.001 using the *S. cerevisiae* proteome as the query against each of the 11 other yeast proteomes independently. We also performed the reciprocal of each of these searches, using each of the 11 other yeast proteomes as the query against the *S. cerevisiae* proteome. We used a custom Python script to identify reciprocal best BLAST hits for each *S. cerevisiae* protein in each of the other yeast proteomes. A protein in the other species’ proteome was considered a reciprocal best hit to the *S. cerevisiae* protein if a) the E-value of the *S. cerevisiae* protein against that protein was the lowest of any in that species’ proteome and b) the E-value of that protein against the *S. cerevisiae* protein was the lowest of any protein in the *S. cerevisiae* proteome. Proteins in the other yeast and insect species satisfying this reciprocal best hit criterion were considered orthologs of the *S. cerevisiae* and protein. When no significant homology to a *S. cerevisiae* protein was detected in another species, or when the reciprocal best hit criterion was not met by any protein in that species, no ortholog was assigned in that species. We repeated this same procedure for all *D. melanogaster* proteins and each of the 21 other insect species’ proteomes.

### Calculation of evolutionary distances

Because evolutionary distance *t* only appears in our model as a product with the gene-specific rate parameter *b*, we can use a subset of genes in the species group to infer these relative distances. Each gene’s value of *b* will scale these relative distances appropriately when fit to the model. We used BUSCO genes as the subset of genes from which to estimate distances, as they are generally well-conserved, facilitating ortholog identification and alignment. We downloaded a list of eukaryotic BUSCO genes [34] from the BUSCO web server (https://busco.ezlab.org/) and identified all of these genes for which we were able to identify an ortholog of the corresponding *S. cerevisiae* gene in all 11 other yeast species (“Identification of orthologs” above). We found 102 such BUSCO genes. We used the alignment software MUSCLE (version 3.8.31) [51] with default parameters to create a multiple sequence alignment of the orthologs from all 12 yeast species for of each of these 102 genes. We then concatenated these alignments and used the Protdist program from the PHYLIP software package (version 3.696) [52] with default parameters to find pairwise evolutionary distances for all 12 yeast species in substitutions per site. To test the effect of using a smaller number of genes to infer these distances, we then randomly and independently selected two subsets of 15 of these 102 genes, and performed the same alignment and distance calculation procedure on each of these two subsets. We then performed the same procedure using *D. melanogaster* genes and the 21 other insect species. Here, there were 125 BUSCOs for which we were able to identify orthologs in all species, and the two random subsets of 15 genes were selected from among these 125. Refseq accessions for genes in the three sets of BUSCOs in both taxa are listed in Supplemental Table 6.

### Correlation of b parameter with evolutionary rate

To determine the correlation between each gene’s best-fit value of the b parameter in our model and the substitution rate, we used alignments of 5261 *S. cerevisiae* genes and their orthologs in all four other *sensu stricto* yeast species generated by a previous study [53]. We opted not to include more distantly related species in these alignments for the sake of more reliable ortholog identification and alignment construction. We used the protdist function of the PHYLIP package (version 3.696) [52] on these alignments to infer the number of substitutions per site between the *S. cerevisiae* gene and its ortholog in the most distant *sensu stricto* yeast *S. kudriavzevii* (we chose a fairly distant representative of these species to minimize sampling error from low substitution counts), and correlated this value with the b parameter inferred from the regression analysis.

### Identification of lineage-specific genes

To identify *S. cerevisiae* genes specific to the three yeast lineages tested here, we performed a BLASTP search [50] with an E-value threshold of 0.001 for each gene in the *S. cerevisiae* proteome as the query against each of the 11 other yeast proteomes independently, using the same proteomes listed in Supplementary Table 2. If the BLASTP search detected no homologs of the *S. cerevisiae* gene in the proteomes of any of these species outside of the specified lineage, we considered it lineage-specific. We applied the same criterion using the 21 other insect proteomes to identify *D. melanogaster* genes specific to the three insect lineages tested here.

### Synteny-based homology searches

We used version 7 of the Yeast Gene Order Browser’s online web tool (http://ygob.ucd.ie/) [40]. For tested *S. cerevisiae* genes, if the gene was included in this YGOB version, we determined whether an orthologous chromosomal region in any of the outgroup yeast species used here had been identified in the browser. If so, we searched for any genes in these outgroup species at the locus that were annotated in the browser. We considered genes to be within the outgroup orthologous locus if they were between the outgroup’s orthologs of the closest *S. cerevisiae* genes up-and downstream of the query gene. If annotated genes existed at the orthologous locus, we performed a BLASTP search of the *S. cerevisiae* sequence against the sequences of all outgroup genes at that locus as listed in YGOB, and called orthology in cases where this single-search E-value was <0.001.

### Gene ontology analysis

We used the Gene Ontology Consortium’s online web server (http://geneontology.org/) [54] to test whether or not certain biological functions were enriched in the set of *sensu stricto*-specific genes that we found to be poorly explained by detection failure. We performed a Fisher’s exact test using the “GO biological process complete” annotation data set for all *S. cerevisiae* genes.

## Supporting information

gzip compressed tar archive of supplementary tables

## Supplemental tables and figures

**Supplemental Table 1a (top), 1b (bottom):**
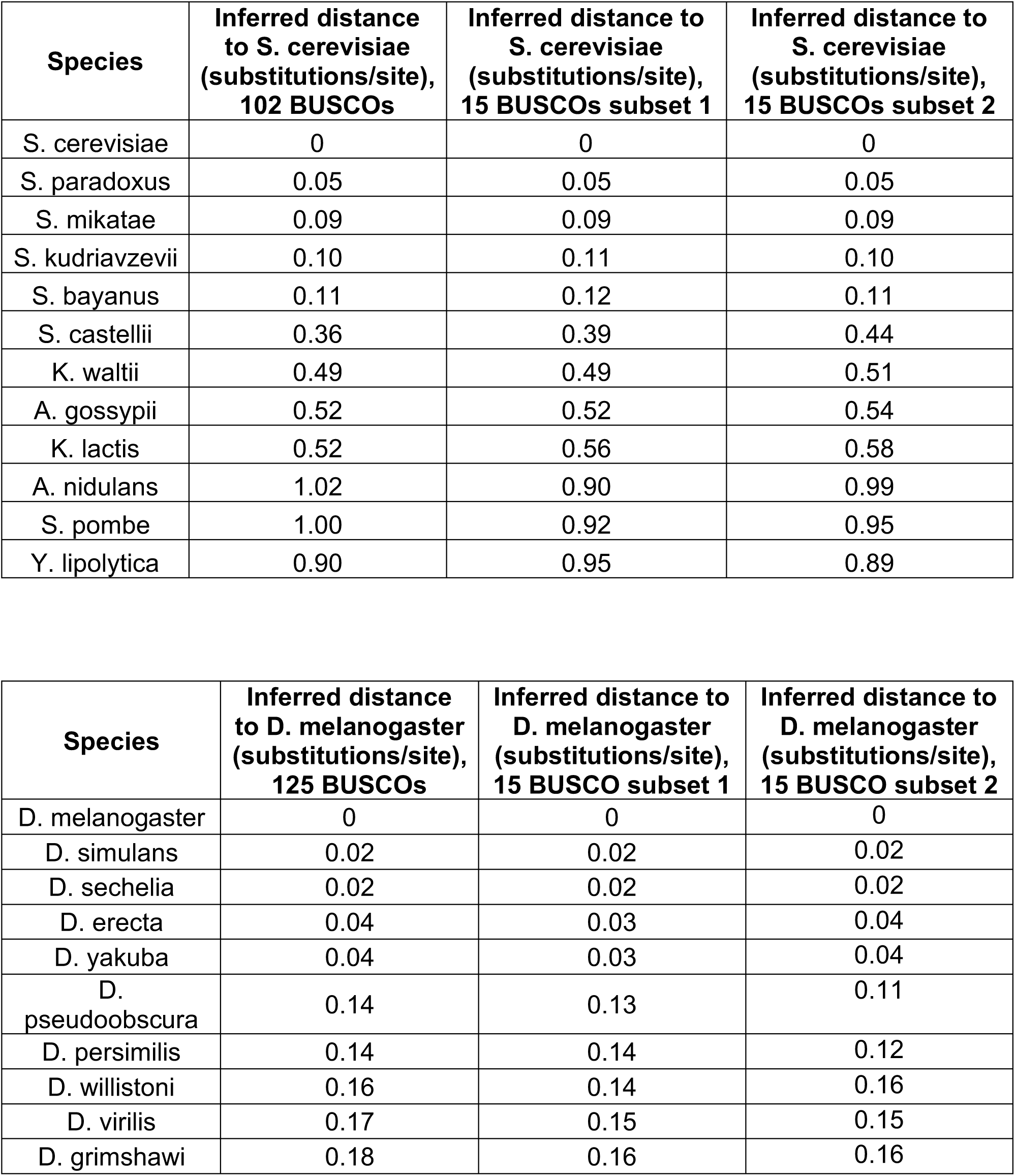
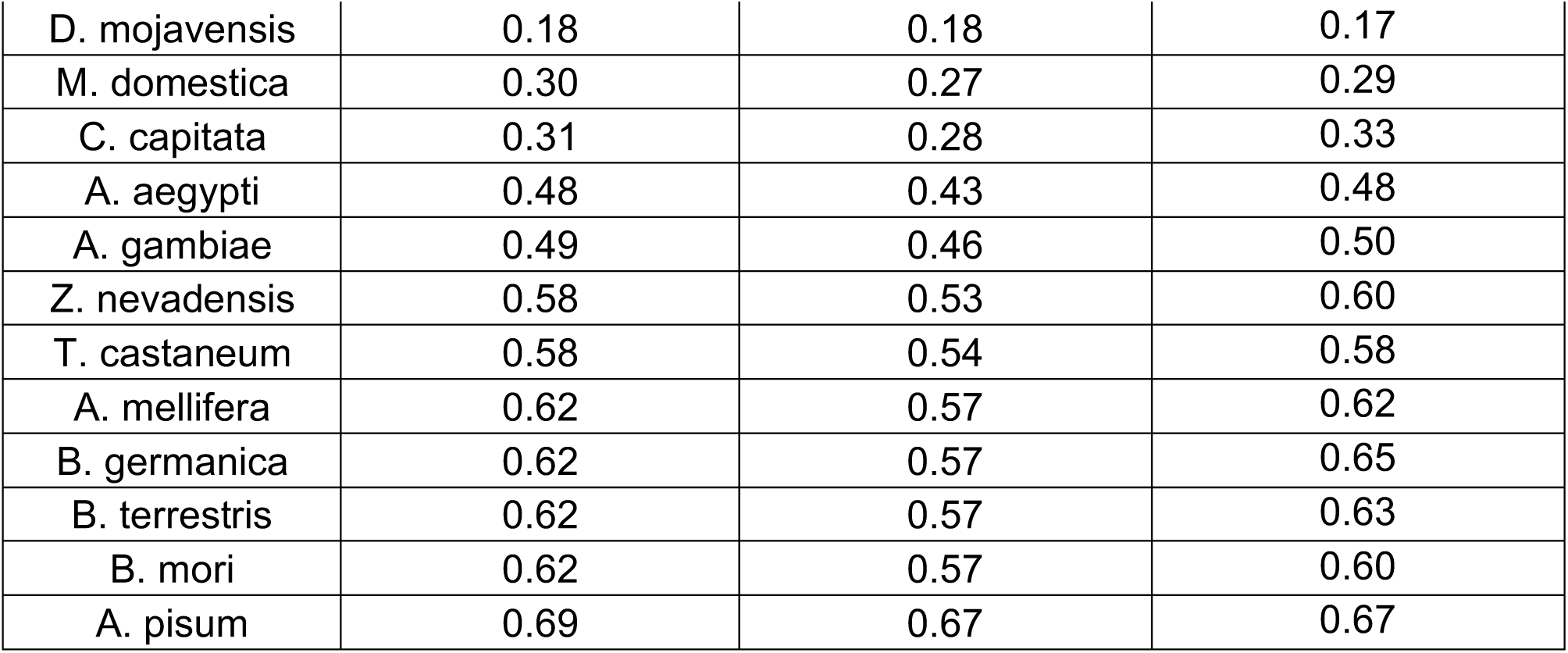
inferred distances in substitutions/site from *S. cerevisiae* to each yeast species (top) and from *D. melanogaster* to each insect species (bottom). Distances were inferred from all BUSCOs with orthologs identifiable in each species group, as well as from 15 genes randomly selected from these BUSCOs. The “15 BUSCOs subset 1” distances were used for all main figures in the text.

| **Species** | **Inferred distance to *S. cerevisiae* (substitutions/site), 102 BUSCOs** | **Inferred distance to *S. cerevisiae* (substitutions/site), 15 BUSCOs subset 1** | **Inferred distance to *S. cerevisiae* (substitutions/site), 15 BUSCOs subset 2** |
| --- | --- | --- | --- |
| *S. cerevisiae* | 0 | 0 | 0 |
| *S. paradoxus* | 0.05 | 0.05 | 0.05 |
| *S. mikatae* | 0.09 | 0.09 | 0.09 |
| *S. kudriavzevii* | 0.10 | 0.11 | 0.10 |
| *S. bayanus* | 0.11 | 0.12 | 0.11 |
| *S. castellii* | 0.36 | 0.39 | 0.44 |
| *K. waltii* | 0.49 | 0.49 | 0.51 |
| *A. gossypii* | 0.52 | 0.52 | 0.54 |
| *K. lactis* | 0.52 | 0.56 | 0.58 |
| *A. nidulans* | 1.02 | 0.90 | 0.99 |
| *S. pombe* | 1.00 | 0.92 | 0.95 |
| *Y. lipolytica* | 0.90 | 0.95 | 0.89 |

Supplemental Table 1a (top), 1b (bottom): inferred distances in substitutions/site from *S.
| **Species** | **Inferred distance to *D. melanogaster* (substitutions/site), 125 BUSCOs** | **Inferred distance to *D. melanogaster* (substitutions/site), 15 BUSCO subset 1** | **Inferred distance to *D. melanogaster* (substitutions/site), 15 BUSCO subset 2** |
| --- | --- | --- | --- |
| *D. melanogaster* | 0 | 0 | 0 |
| *D. simulans* | 0.02 | 0.02 | 0.02 |
| *D. sechelia* | 0.02 | 0.02 | 0.02 |
| *D. erecta* | 0.04 | 0.03 | 0.04 |
| *D. yakuba* | 0.04 | 0.03 | 0.04 |
| *D. pseudoobscura* | 0.14 | 0.13 | 0.11 |
| *D. persimilis* | 0.14 | 0.14 | 0.12 |
| *D. willistoni* | 0.16 | 0.14 | 0.16 |
| *D. virilis* | 0.17 | 0.15 | 0.15 |
| *D. grimshawi* | 0.18 | 0.16 | 0.16 |
| D. mojavensis | 0.18 | 0.18 | 0.17 |
| M. domestica | 0.30 | 0.27 | 0.29 |
| C. capitata | 0.31 | 0.28 | 0.33 |
| A. aegypti | 0.48 | 0.43 | 0.48 |
| A. gambiae | 0.49 | 0.46 | 0.50 |
| Z. nevadensis | 0.58 | 0.53 | 0.60 |
| T. castaneum | 0.58 | 0.54 | 0.58 |
| A. mellifera | 0.62 | 0.57 | 0.62 |
| B. germanica | 0.62 | 0.57 | 0.65 |
| B. terrestris | 0.62 | 0.57 | 0.63 |
| B. mori | 0.62 | 0.57 | 0.60 |
| A. pisum | 0.69 | 0.67 | 0.67 |

Supplemental Table 2 (attached): Sources of species protein annotations used in this study.

Supplemental Table 3 (attached): Correlation coefficients for gene detectability prediction results based on evolutionary distance estimates derived from three different sets of genes (the same as those shown in Supplemental Table 1).

Supplemental Table 4 (attached): List of 11 *S. cerevisiae* genes for which synteny-based searches in YGOB revealed candidate out-of-lineage orthologs, the YGOB IDs of those orthologs, and their synteny search E-values.

Supplemental Table 5 (attached): List of *sensu stricto*-specific *S. cerevisiae* genes that are poorly explained by the hypothesis of detection failure and their features as described in summary in the text. (Attached file.)

Supplemental Table 6 (attached): List of RefSeq accession IDs for BUSCOs used in evolutionary distance calculations.

**Supplemental Figure 1a-f:**
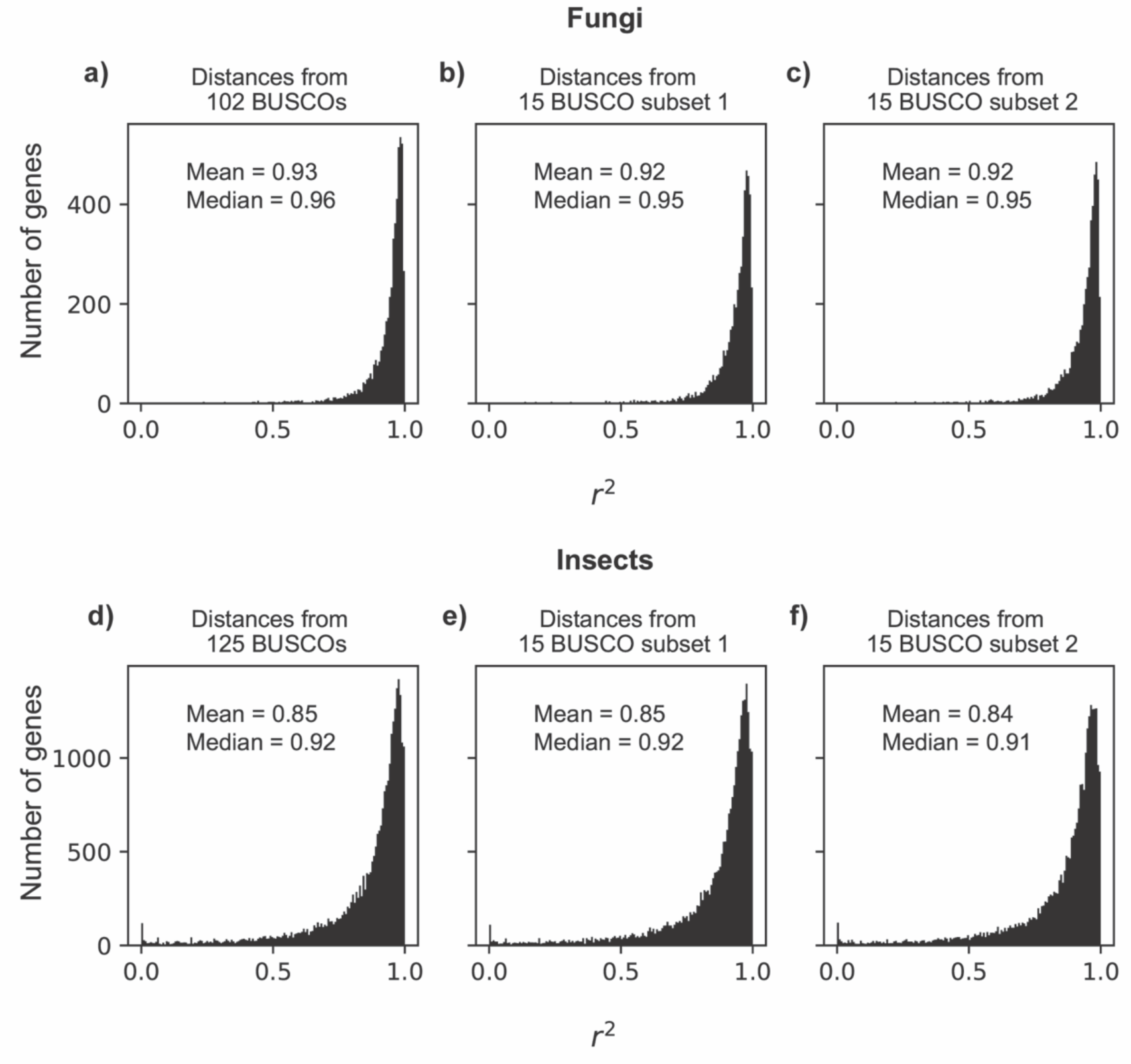
r^2^ distributions for the fit to the model of *S. cerevisiae* and *D. melanogaster* genes using evolutionary distances derived from three sets of genes. **a**: *S. cerevisiae* genes with distances derived from 102 BUSCOs. **b**: *S. cerevisiae* genes with distances derived from a randomly-selected subset of 15 of the BUSCOs used in a. **c**: *S. cerevisiae* genes with distances derived from a second randomly-selected subset of 15 of the BUSCOs used in a. **d**: *D. melanogaster* genes with distances derived from 125 BUSCOs. **e**: *D. melanogaster* genes with distances derived from a randomly-selected subset of 15 of the BUSCOs used in d. **f**: *D. melanogaster* genes with distances derived from a second randomly-selected subset of 15 of the BUSCOs used in d.

**Supplemental Figure 2:**
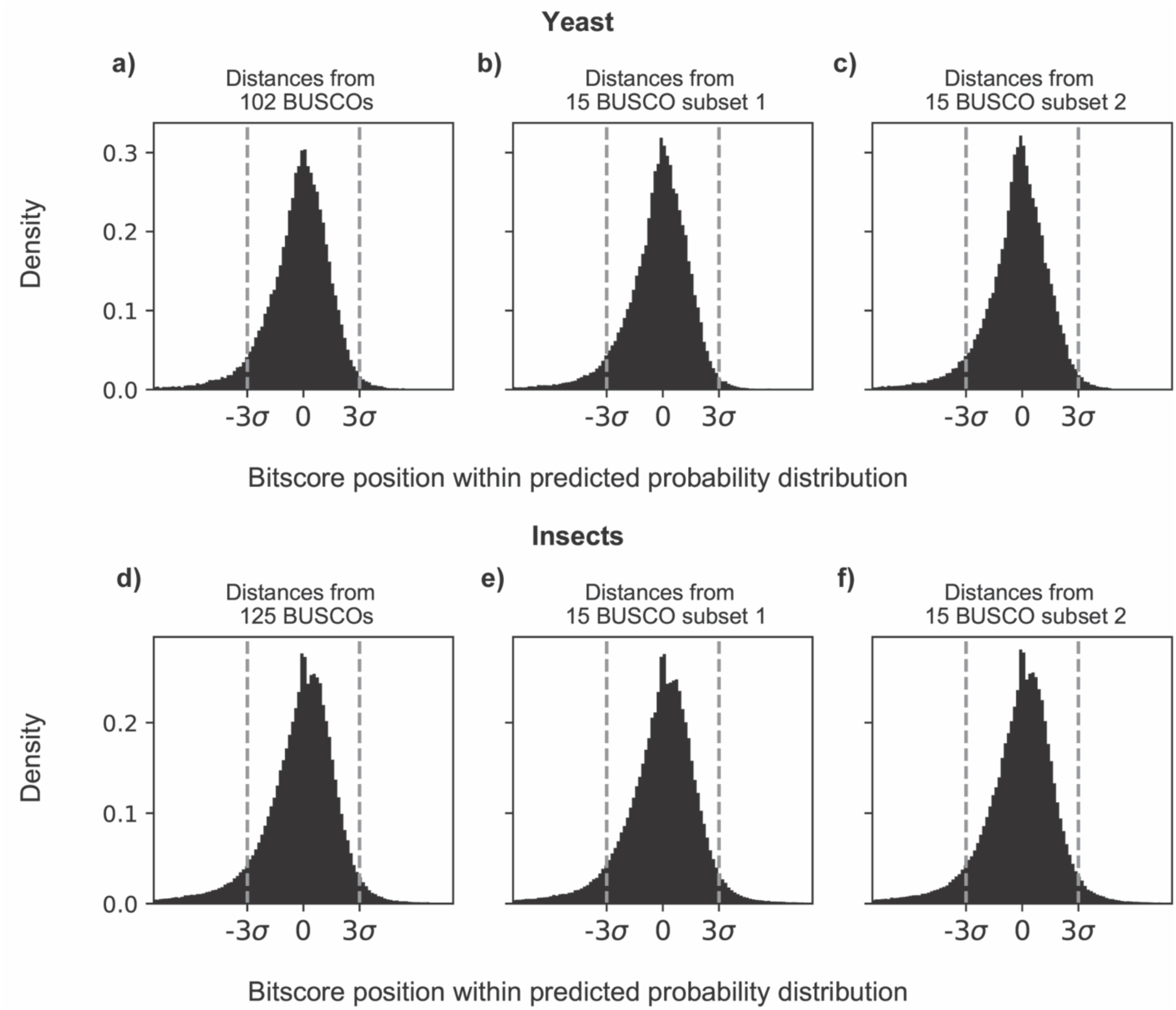
Distribution of position of BLASTP scores between *S. cerevisiae* and outgroup yeast (top) and *D. melanogaster* and outgroup insects (bottom) relative to the predicted confidence interval. 0 indicates that the score has the same value as the best fit to the model; multiples of sigma indicate that the score is that many standard deviations above or below the best fit value. **a**: *S. cerevisiae* genes with distances derived from 102 BUSCOs. **b**: *S. cerevisiae* genes with distances derived from a randomly-selected subset of 15 of the BUSCOs used in a. **c**: *S. cerevisiae* genes with distances derived from a second randomly-selected subset of 15 of the BUSCOs used in a. **d**: *D. melanogaster* genes with distances derived from 125 BUSCOs. **e**: *D. melanogaster* genes with distances derived from a randomly-selected subset of 15 of the BUSCOs used in d. **f**: *D. melanogaster* genes with distances derived from a second randomly-selected subset of 15 of the BUSCOs used in d.

**Supplemental figure 3a,b:**
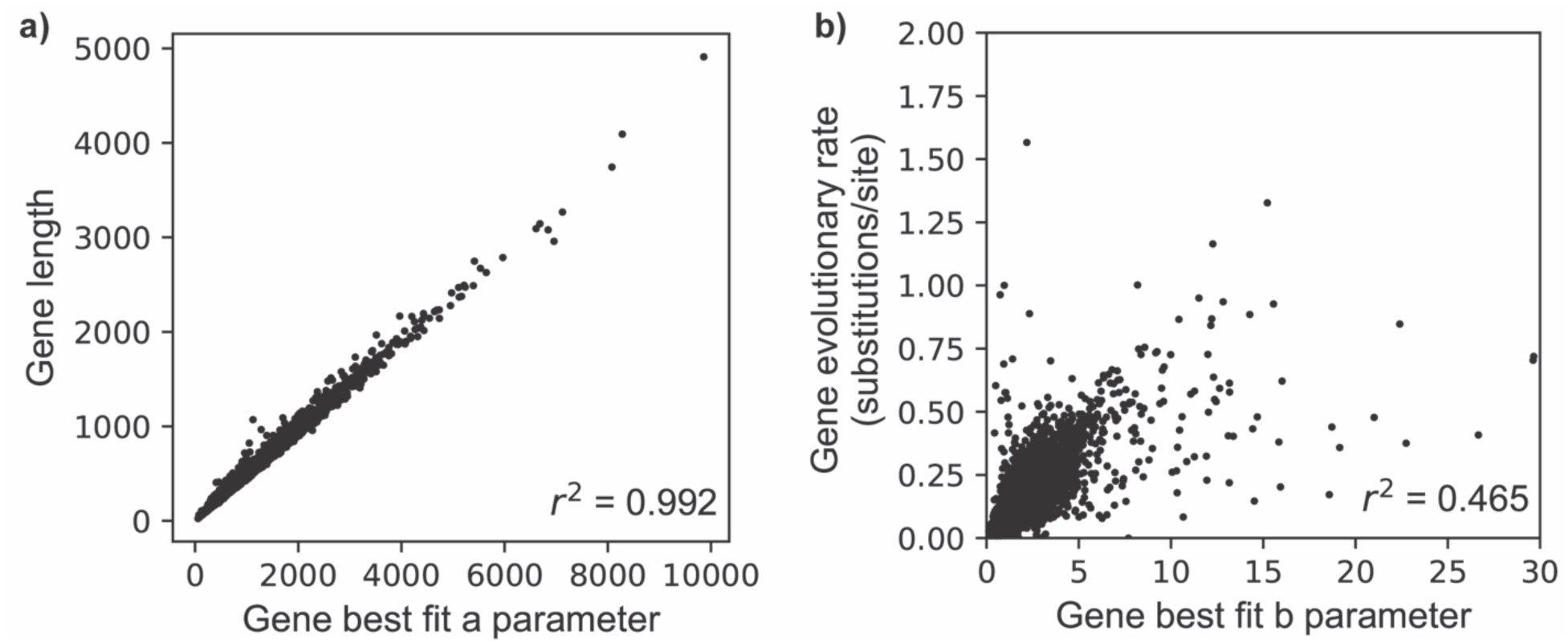
Correlation between best-fit parameters and gene properties in yeast. 3a: Correlation between each *S. cerevisiae* protein’s best-fit value of *a* and its length in amino acids. The *a* parameter is consistently larger than the length due to most identical alignment positions contributing a score larger than 1 according to the scoring scheme used here (BLOSUM62). 3b: Correlation between each *S. cerevisiae* protein’s best-fit value of *b* and its relative evolutionary rate in substitutions per site from *sensu stricto* protein alignments (Methods).

**Supplemental figure 4a,b:**
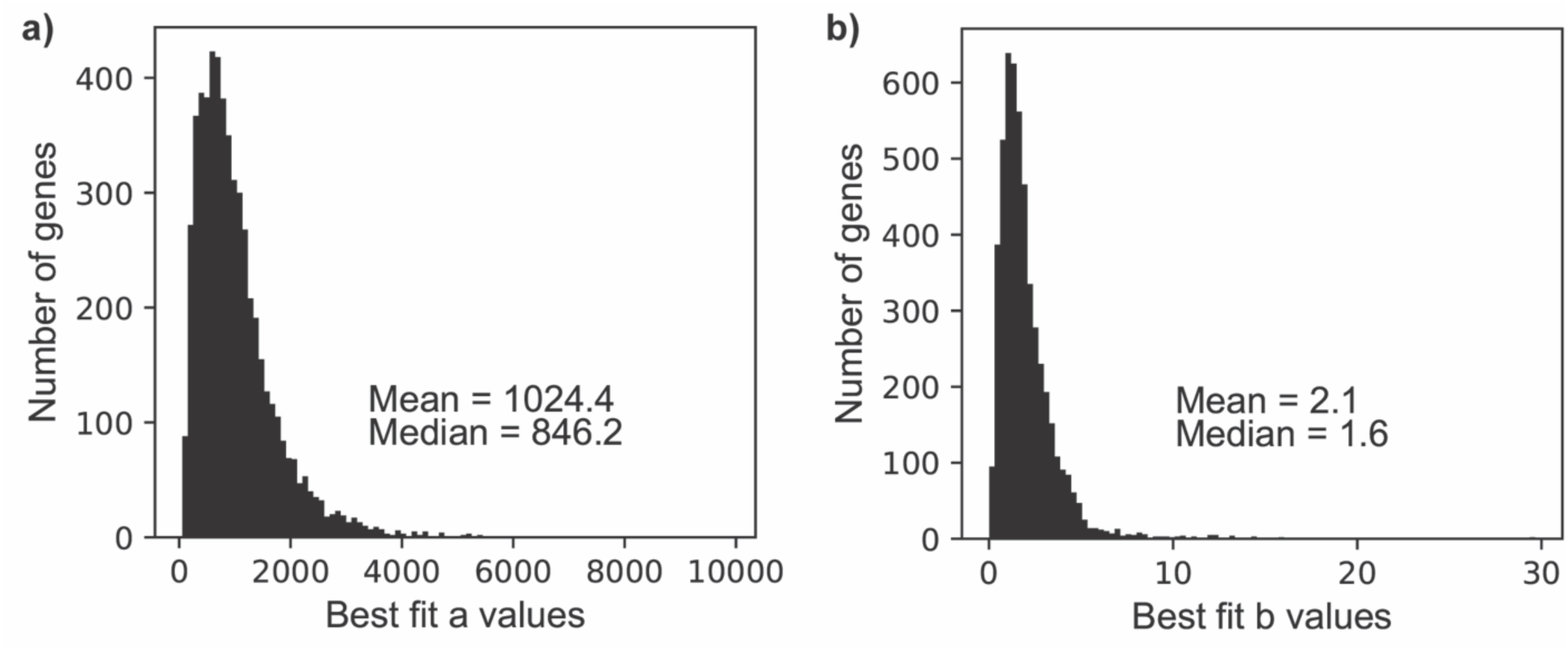
Distribution of best fit parameter values for all *S. cerevisiae* proteins. 4a: Distribution of the best-fit *a* values for all *S. cerevisiae* proteins. 4b: Distribution of the best-fit *b* values for all *S. cerevisiae* proteins.

